# Chimeric aggregative multicellularity in absence of kin discrimination

**DOI:** 10.1101/2024.12.04.626738

**Authors:** Michael L. Weltzer, Jack Govaerts, Daniel Wall

## Abstract

Aggregative multicellularity is a cooperative strategy employed by some microorganisms. Unlike clonal expansion within protected environments during multicellular eukaryotic development, an aggregation strategy introduces the potential for genetic conflicts and exploitation by cheaters, threatening the stability of the social system. *Myxococcus xanthus*, a soil-dwelling bacterium, employs aggregative multicellularity to form multicellular fruiting bodies that produce spores in response to starvation. Studies of natural fruiting bodies show that this process is restricted to close kin or clonemates. Here, we investigate the mechanisms underlying kin recognition during development in *M. xanthus*. By co-culturing two distantly related *M. xanthus* strains under vegetative and starvation conditions, we observed that the strains segregate in both contexts. During vegetative growth, one strain antagonized the other using the type VI secretion system (T6SS). T6SS-mediated antagonism was also observed during development, resulting in monoclonal fruiting bodies when WT strains were mixed. In contrast, mixtures of T6SS knockout strains formed chimeric fruiting bodies, that produced viable spores from both strains. These findings suggest that T6SS is the primary mechanism of kin discrimination in distantly related *M. xanthus* strains, and its use ensures the development of monoclonal fruiting bodies and social integrity.

## Introduction

Multicellularity requires cells to cooperate to form functional tissues and for individuals to reach maturity. Failure to cooperate leads to disease, such as cancer, or non-viability [1]. Plants and animals create unique genetic offspring by gamete fusion to form a single zygotic cell from which all subsequence cells are derived. This single cell bottleneck serves as a checkpoint to ensure all cells are genetically identical and as a purifying mechanism to remove genotypes that hinder cooperation. In contrast, some species use an aggregation strategy where cells coalesce from their environment to build a multicellular organism. This latter strategy is ripe for genetic conflict between non-clonal cells. This includes exploitation by cheaters, cells that utilize resources from cooperative communities without contributing their fair share of public goods [2, 3].

For the aggregation strategy to succeed, cells evolved mechanisms to distinguish self from nonself. This occurs by recognition and/or discrimination mechanisms. As defined here, kin recognition refers to cells that use genetic determinants to identify other cells that are clonal or highly related to conduct cooperative and beneficial acts. In contrast, kin discrimination refers to cells that conduct antagonistic acts toward cells that are not close kin. Two model systems used for understanding self/nonself recognition during aggregative multicellularity are the eukaryotic social slime mold *Dictyostelium discoideum* and the social bacterium *Myxococcus xanthus*. These extremely divergent species share lifestyles where they are soil microbial predators and in response to starvation, thousands of cells aggregate to build fruiting bodies wherein cells differentiate into different types including stress resistant spores. In the case of *D. discoideum*, they primarily use kin recognition that involves heterotypic binding between polymorphic cell surface adhesins called TgrB1 and TgrC1 [4]. Cell-cell binding, mediated by these proteins, triggers actin cytoskeleton remodeling and motility driven segregation between strains with different allotypes or alleles of *tgrB1/C1*. Strain segregation occurs early during the initial aggregation of cells as well as later developmental stages and does not involve direct antagonism or killing. Notably, TgrB1/C1 allorecognition also protects cooperative populations from cheaters cells [5].

*M. xanthus* also uses kin recognition mediated by a polymorphic cell surface receptor called TraA and its cohort protein TraB. Self-recognition occurs by homotypic binding where specificity is determined by TraA polymorphisms [6–8]. This leads to the bidirectional exchange of outer membrane proteins and lipids, called outer membrane exchange (OME), which can endow benefits to kin [9, 10]. However, to form homogenous developmental populations from their diverse environments, we hypothesize this is primarily driven by kin discrimination, because prior studies showed conspecific strains intensely antagonize one another thus presumably precluding multicellular development [11]. Kin discrimination mechanisms include OME that delivers dozens of different polymorphic toxins to divergent neighboring cells that happen to express a compatible *traA* allele [12, 13]. In contrast, clonal cells are protected because they express a cognate suite of immunity proteins that are not transferred. A second and broader kin discrimination system involves polymorphic toxin delivery by the type VI secretion system (T6SS), an injection platform evolutionarily related to phage tails [14]. Apparently, the T6SS injects toxins in a nonspecific manner into neighboring cells and if clonal, they similarly express the cognate set of immunity proteins. Curiously, our lab and others found that *M. xanthus* does not use T6SS for bacterial predation, but instead it serves as a major kin discrimination determinant against other myxobacteria [11][15, 16].

During starvation, over 10^5^ *M. xanthus* cells aggregate to form a fruiting body. However, only 5-20% of cells differentiate into spores [17], while other cells become peripheral rods (a.k.a. persisters) but the majority, approximately 80%, lyse [18][19]. Because of this, fruiting bodies are vulnerable to exploitation by cheaters that do not lyse. Indeed, this has been observed under laboratory conditions in which developmentally deficient lines evolved under asocial conditions were overrepresented as spores when mixed with their unevolved ancestor [2]. Cheating has severe consequences, including drastically altering population dynamics or causing complete collapse of the social system and population extinction [20]. Importantly, fruiting bodies from the wild are composed of nearly genetically identical individuals [21] indicating that *M. xanthus* has mechanisms to determine genetic relatedness and exclude non-kin cells, which could be cheaters.

Previously, we investigated kin discrimination between pairs of closely related natural isolates growing vegetatively [11]. Here, *M. xanthus* was isolated from a 16 cm × 16 cm grid of forest soil [22][23], where pairs of isolates were grown and spotted next to one another on agar surfaces and grouped into compatibility types based onto whether the swarm colonies merged or not. In some cases, isolates that were nearly genetically identical, were incompatible [24][23]. We compared the draft genomes of these isolates and found that they contained different genomic islands, primarily of prophage origin, which carry unique *sitA* toxin loci (delivered by OME) and T6SS toxin loci. We found that when mixed, wild-type strains antagonize one another and genetically inactivating both OME and T6SS ceased the killing behaviors. However, for some strains, double knockouts mutants continued to antagonize others. These strains each contained a large and unique *rhs* toxin genes in prophage islands and knocking out an *rhs* gene, in addition to the two other toxin delivery systems, relieved antagonism [11].

In this work, we asked if preventing antagonism in *M. xanthus* is sufficient to allow non-kin cells to engage in cooperative behaviors, such as swarming and fruiting body formation. To do so we disabled known kin discrimination mechanisms and tested whether divergent *M. xanthus* strains harmoniously coexisted, or if other kin recognition or discrimination mechanisms are involved. We tested these questions using two distantly related *M. xanthus* strains: Environmental isolate A06 from a German forest and the lab strain DK1622 originally isolated from Ames Iowa. These isolates contain incompatible *traA* alleles and therefore cannot engage in OME. Thus, T6SS mutants, labelled with fluorescent proteins were found not to antagonize one another under vegetative conditions and can merge to form chimeric fruiting bodies that produce viable spores in response to starvation. These compatible interactions were compared with the incompatible interactions between the parent strains and the implications of these findings are discussed.

## Materials and Methods

### Strain construction

To create a labeled strain of DK1622, pMW106 (tdTomato and oxytetracycline resistance) was electroporated and recombined into the genome of DK1622 (WT). To create labeled strains of A06, either plasmid pMW106 or pMW119 (GFP and streptomycin resistance) was transformed and integrated into the A06 genome or similarly to a previously created T6SS KO mutant of A06 [11]. Transformants were selected on CTT agar media (1% Casitone, 10 mM Tris-HCl [pH 8.0], 1 mM K2HPO4, 8 mM MgSO4, [final pH 7.6]), supplemented with either 50 μg/mL kanamycin, 10 μg/mL oxytetracycline or 1.5 mg/mL streptomycin.

### Growth and development

Strains were grown overnight to mid log phase in CTT. For experiments under vegetative conditions, strains were resuspended to a density of 7.5 X 10^8^ cells/mL, mixed at a 1:1 ratio, and spotted on CTT 1% agar. At each time point, spots were scraped, resuspended in liquid TPM and cells were enumerated using fluorescence microscopy (see below). For starvation experiments on an agar surface, cells were harvested and resuspended to a density of 3 X 10^9^ cells/mL on TPM (10 mM Tris-HCl [pH 8.0], 1 mM K_2_HPO_4_, 8 mM MgSO_4_, [final pH 7.6]) 1% agar plates. For submerged culture, strains were mixed at a 1:1 ratio at an initial cell density of 3 X10^7^ cells/mL and grown in 500 μL of CTT for 24 h in a 24-well plate. At 24 h, CTT was removed and 1mL of MC7 buffer (10 mM morpholinepropanesulfonic acid [pH 7.0] and 1 mM CaCl_2_) was added.

### Sequencing and phylogenetic analysis

For sequencing, strains A06 and DW2653, a derivative of A44, were grown overnight and genomic DNA was harvested using the Wizard Genomic DNA Purification Kit (Promega). Nanopore sequencing and genome annotation was performed by SeqCenter (Pittsburgh, PA).

For phylogenetic analysis, eight fully sequenced publicly available *M. xanthus* genomes were obtained from IMG. From these eight genomes, DK1622, A06 and A44 we performed MLST analysis using seven housekeeping genes: *dnaA*, *gyrB, pyrG, rpoB, lepA, fusA,* and *secA.* 60 nucleotides were missing from the beginning of some of the *dnaA* sequences, so these nucleotides were removed from the analysis. Sequence*s* were aligned with Clustal Omega [25, 26]. The aligned sequences were analyzed by ModelTest-NG [27] and the TIM2 +I +Gamma model of DNA substitution was selected. A maximum likelihood phylogeny was created with RAxML-NG v1.1.0 [28]using this DNA substitution model and 10,000 transfer bootstrap expectation replicates.

### Microscopy

For high magnification images of swarms, cells were spotted onto 1% agar CTT pads and imaged using an Olympus IX83 inverted microscope (40× lens objective coupled to an ORCA-Flash 4.0 LT sCMOS camera and cellSens software). Low magnification images of swarms and fruiting bodies were captured using an Olympus SZX10 stereomicroscope coupled to a digital imaging system. Fluorescent images of fruiting bodies were captured with a Nikon E800 microscope (2× or 10× lens objective coupled to an ORCA-Flash 4.0 LT sCMOS camera and cellSens software).

### Spore assay

Strains were grown overnight in CTT at 33 °C to early log phase, then resuspended to a density of 3 X10^9^ cells/mL in liquid TPM. 10 μL of monocultures or 20 μL of a 1:1 mixture of strains was spotted on TPM 1% agar plates. After 5 days of development, six spots were scraped from the plate and resuspended in TPM. The suspension was heat treated at 50 °C for 2 h to kill vegetative cells and sonicated to break apart spore clumps. Sonication was performed on ice 3 times for 30 s each with 20 s between each sonication. Following sonication, spores were serially diluted and plated on either CTT supplemented with 2.5 μg/mL oxytetracycline or 500 μg/mL streptomycin.

## Results

### Genomic comparison of DK1622 and A06

In this study, we investigated the interactions between two *M. xanthus* strains that were isolated on different continents, decades apart: environmental isolate A06 and the well-studied lab strain DK1622. To determine the evolutionary relationship between these and other *M. xanthus* strains, we conducted multilocus sequence typing of seven conserved housekeeping genes (Fig. 1A).

**Fig. 1.**
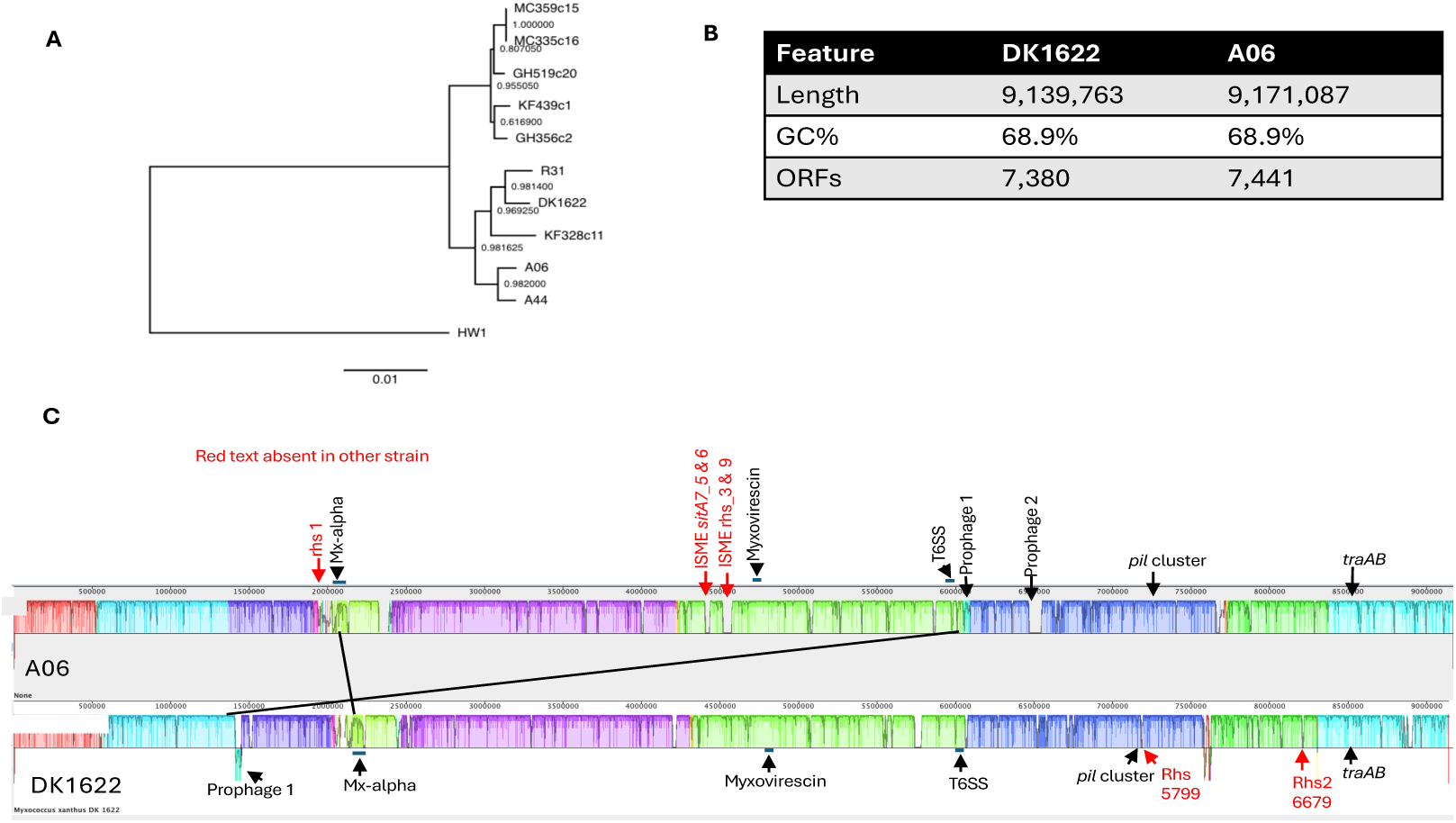
Phylogenic relationship between A06 and DK1622. **A.** Maximum likelihood phylogeny showing the relationship of *M. xanthus* strains using seven conserved housekeeping genes: *dnaA, gyrB, pyrG, rpoB, lep, fusA* and *secA.* Numbers at nodes represent bootstrap values. Outgroup is *Myxococcus macrosporus* HW1. **B.** Genome comparison between strains. **C.** Alignment of A06 and DK1622 genomes using progressiveMauve [29]. Areas of the same color represent homologous regions. Selective elements and genes indicated. Genes in red text are unique to one strain.

A06 grouped in a clade with A44, an isolate from the same soil patch, which shared 99.43% DNA sequence identity across the assembled 16,286 bp fragment. DK1622 grouped in a sister clade to these isolates and shared 98.63% DNA sequence identity with A06.

Next, we used comparative genomics to further understand the genetic relationship between these strains. The DK1622 genome was previously sequenced [30] and we used nanopore technology to sequence the A06 genome. As expected, the 16S rRNA genes were 99.93% identical (single base difference). We aligned the two genomes with Mauve and found overall, their genomes had close colinearity or synteny, including the large genomic islands of prophage Mx-alpha and the myxovirescin polyketide biosynthetic gene cluster (Fig 1C), which are found in some but not all *M. xanthus* genomes. In contrast, some islands were missing in one strain or located in a different region. We additionally compared genes encoding cell surface products, which are frequently involved in cell-cell interactions and are more prone to sequence polymorphisms. In the case of *t6ss*, *eps* and *csgA* genes, which function as transport systems or enzymes, they showed a high degree of similarity, e.g. 98 to 100% protein sequence identify (Fig 2). In contrast, genes that encode cell surface recognition proteins, including *traA, pilA,* and *pilY1.1,* were divergent, e.g. sharing 78.86% to 82.25% protein sequence identify, suggesting these strains belong to different social groups. Nearly all the sequence differences in *traA* were in the variable domain involved in kin recognition specificity [6, 8, 31]. Consequently, these strains are incompatible for OME and kin discrimination mediated by the transfer of polymorphic SitA toxin families.

**Fig. 2.**
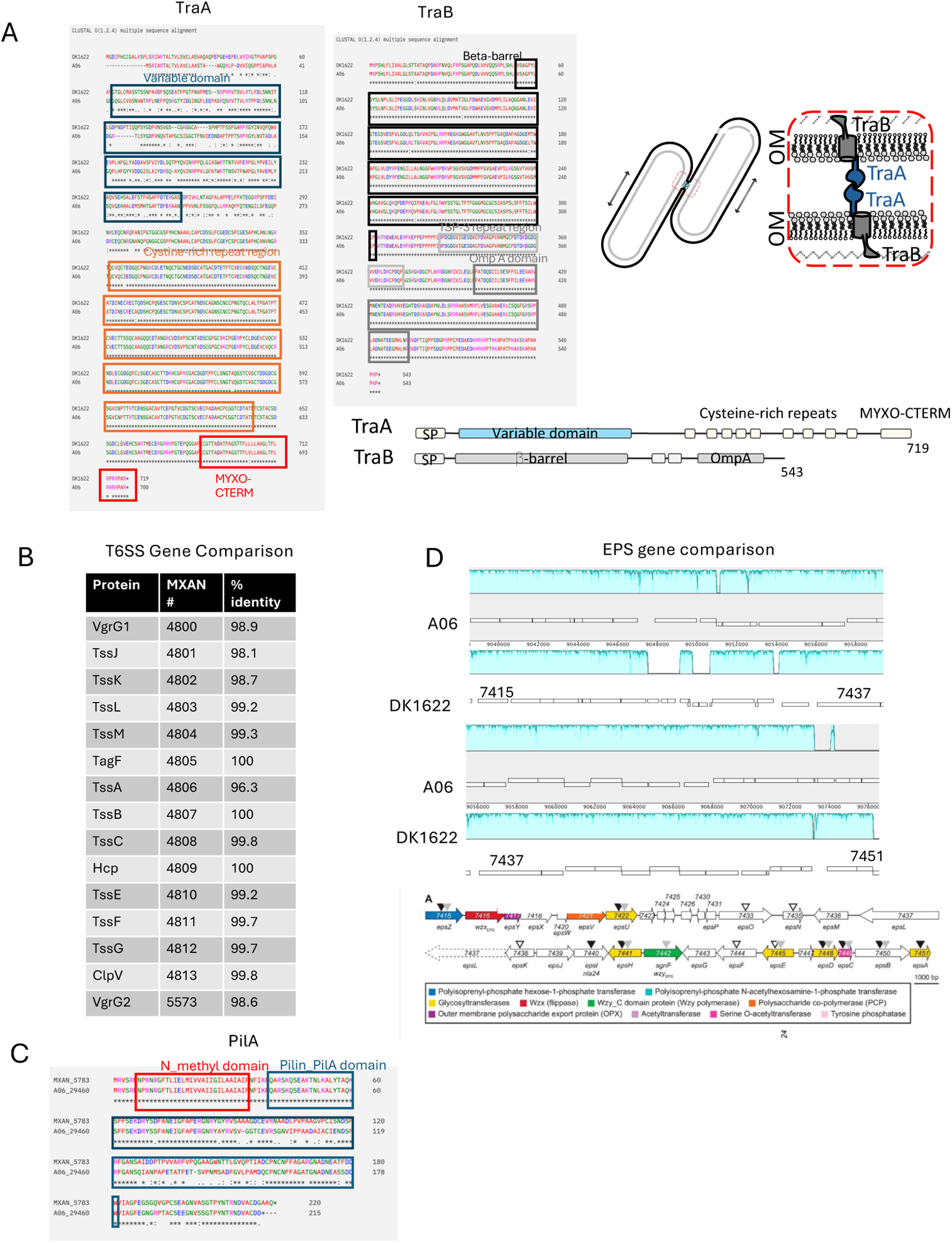
Comparison of social genes between A06 and DK1622. **A.** Sequence alignment of TraA and TraB proteins. Known protein domains boxed. Cartoon depicts model of homotypic TraA-TraA binding, between compatible TraA receptors. **B.** Percent identity of T6SS proteins. MXAN# locus tags of DK1622 shown. **C.** Sequence alignment of PilA proteins. Known domains boxed. **D.** Mauve alignment of EPS gene clusters (upper). Colored regions indicate homology. Numbers indicate MXAN locus tags from DK1622. Description of EPS ORFs in DK1622 adapted from [32](lower).

Our prior work revealed that polymorphic effectors involved in kin discrimination are shuffled in diverse combinations in strains apparently driven by horizontal gene transfer [11]. We then compared toxin genes in A06 to DK1622. As expected, each genome contained unique sets of SitA (Table 1A) and T6SS toxin genes (Table 1B). Of the 34 total *sitA* genes found, only three sets showed significant identity (97 to 100%) between strains, suggesting their cognate immunity genes provide cross-resistance. For T6SS effectors, of the 13 total identified, only three showed significant identity between strains (98.9% to 99.7%), again suggesting cross-resistance to those effectors by their cognate immunity factors. Some of these toxin-immunity cassettes were found on large prophage elements, described previously [11]. We performed BLAST searches on T6SS toxins unique to A06 and found that they had close homologs in other myxobacteria (Table 1C). Additionally, each strain had unique *rhs* genes that might contribute toward kin discrimination (Fig. 1C). Nevertheless, based on the different sets of T6SS effectors, we predicted these strains would antagonize each other via their T6SS.

**Table 1.**
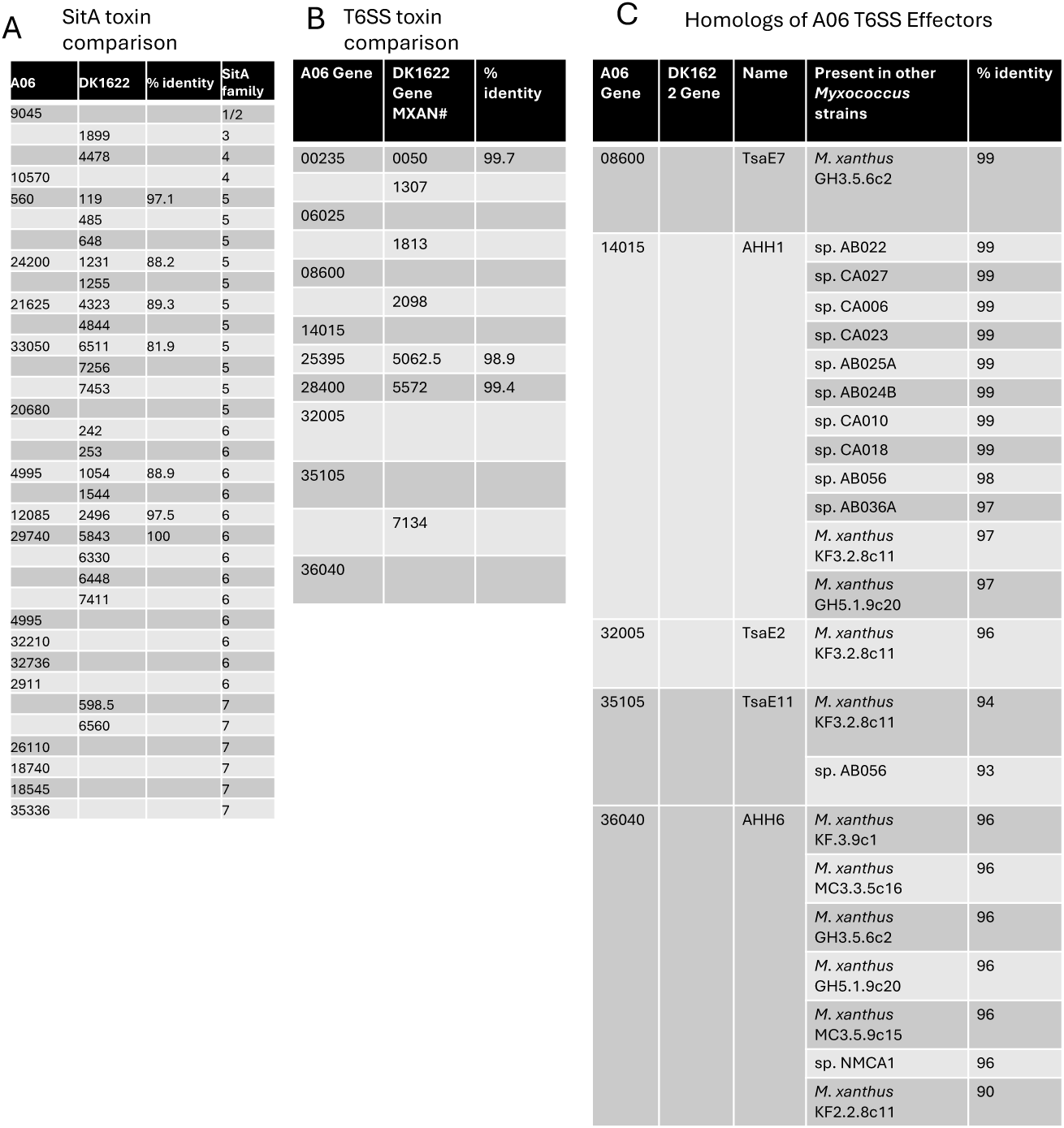
Sequence comparisons of OME and T6SS effectors between A06 and DK1622. (A) Amino acid sequence identities between SitA toxins when homologs are present between strains. (B) Amino acid sequence identities between T6SS toxins when homologs are present between strains. (C). BLAST hits and protein identities of A06 T6SS toxins from other myxobacteria. Sequences ≥ 95% identical are considered the same allele.

### During vegetative growth, T6SS antagonism occurs

To test for inter-strain antagonism, we spotted A06 and DK1622 next to one another on rich media agar plates. As has previously observed for distant strains [24], a clear demarcation formed between the swarms, while swarms between clonal colonies freely merged. To test for the mechanism of antagonism, we constructed T6SS mutants and placed them next to one another. In this pairing, no demarcation emerged (Fig 3A).

**Fig. 3.**
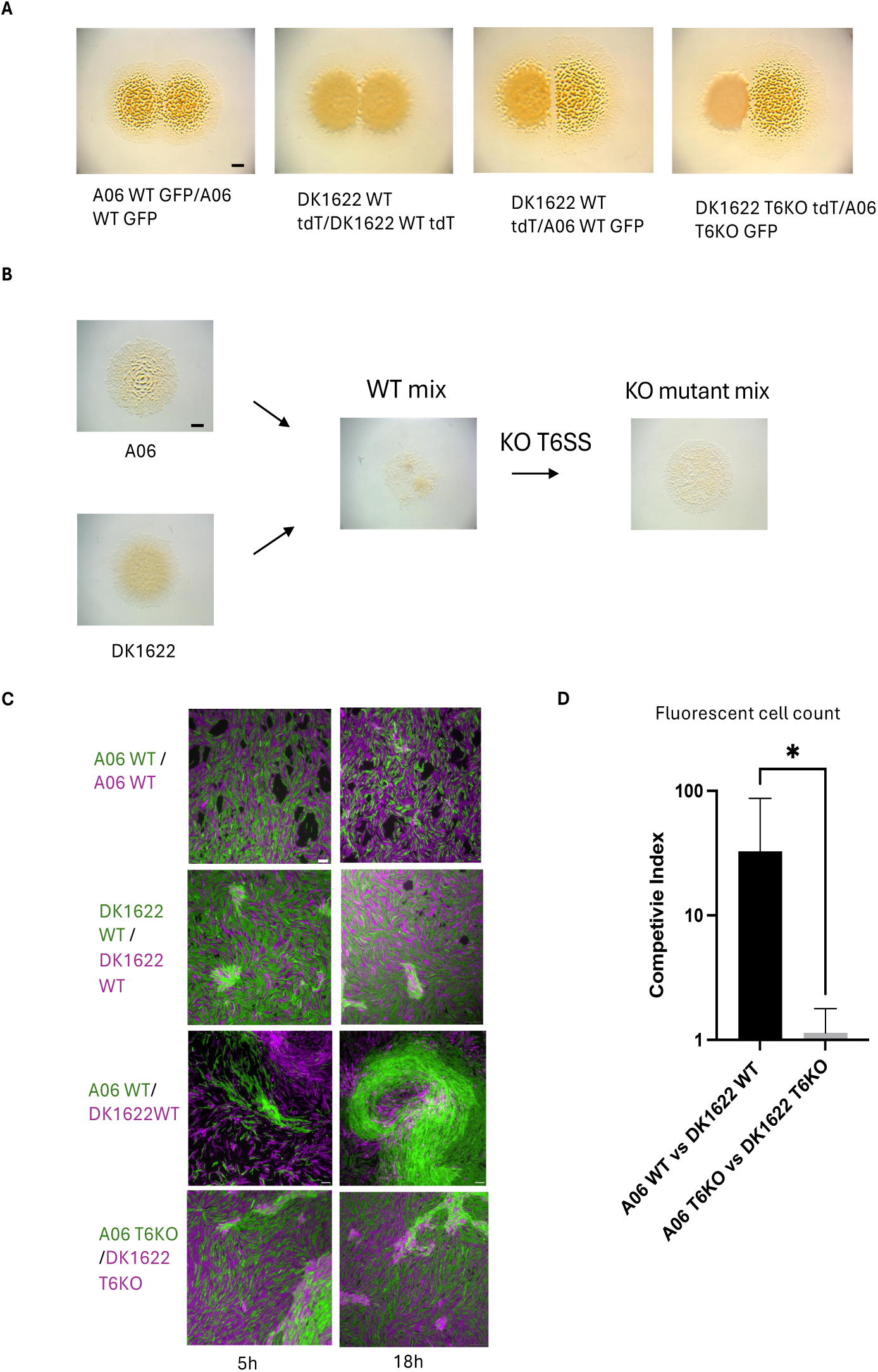
Social interactions between A06 and DK1622 derived strains. **A.** Colony-merger incompatibility test. Aliquots of each strain spotted next to one another on CTT 1% agar plates. Micrographs taken at 48 h. Scale bar, 1 mm. **B.** Aliquots of monocultures or 1:1 mixtures of indicated strains on CTT agar. Micrographs taken at 24 h. Scale bar, 1000 µm. **C.** Fluorescent micrographs of 1:1 strain mixtures taken at indicated times after spotting on 1% CTT agar pads. A06 and DK1622 derived strains labeled with GFP or mCherry, respectively. Scale bar, 10 µm. **D.** Quantification of competition experiments of 1:1 mixtures of WT or T6SS KO strains. Mixtures were spotted on CTT plates, collected at 24 h and number of cells from each strain were enumerated by fluorescence microscopy determined their competitive index (ratio between strains). *P < 0.05 unpaired t-test.

Next, we mixed the WT strains together at a 1:1 ratio and transferred them on rich agar media, as well as respective monoculture controls. 24 h after spotting, we observed sparse growth of the mixture, which contrasted with robust monoculture growth (Fig 3B), further indicating inter-strain antagonism. However, when corresponding T6SS mutants were mixed, robust growth occurred to near monoculture levels, indicating antagonism was greatly reduced or eliminated. To quantify the level of antagonism, we labeled each strain with different fluorescent and antibiotic resistance markers. DK1622 was labeled with tdTomato and tetracycline resistance, while A06 was labeled with GFP and kanamycin resistance. Strains were mixed 1:1 and monitored by fluorescent microscopy. At 5 h post mixing cell debris from both strains was readily seen (Fig 3C) revealing mutual antagonism. By 18 or 24 h, nearly all the DK1622 cells were absent (Fig. 3C-D); indicating A06 was the dominant strain. As controls, isogenic strains were mixed expressing the two different fluorescent markers and no antagonism was detected for DK1622 and A06 (Fig. 3.3C) at 5 or 18 h.

To assess the role of T6SS in antagonism, we similarly mixed T6SS mutants, with the same markers. At 5 and 18 h, we found no evidence of antagonism, and the strain ratio remained unchanged at 24 h (Fig. 3D). We conclude that under vegetative growth, A06 and DK1622 antagonism was primarily or solely mediated by their T6SS and A06 was the dominant competitor.

### Chimeric fruiting bodies form in the absence of T6SS antagonism

To investigate strain interactions during development, we placed a 1:1 mixture of WT strains on starvation agar and monitored development as compared to monoculture controls (Fig. 4A). Strikingly, few mature fruiting bodies emerged after 5 days, which contrasted with monoculture development. Importantly, the mixture of T6SS mutants restored fruiting body development (Fig. 4A).

**Fig. 4.**
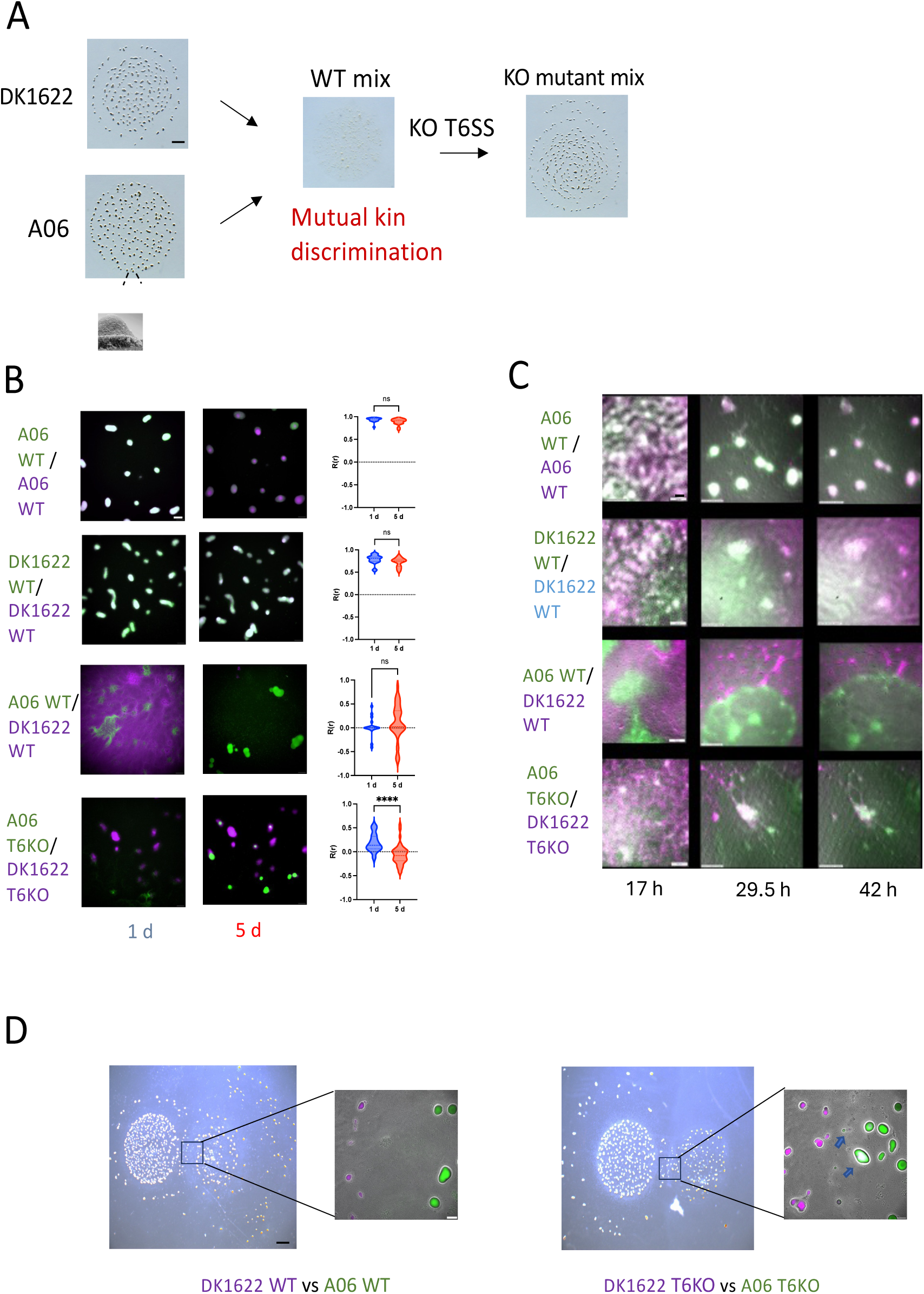
Developmental interactions between A06 and DK1622 derived strains. **A.** Aliquots of monocultures or 1:1 strain mixtures on TPM 1% agar plates at 72 h. Dark spots are fruiting bodies (arrow). Scale bar, 1 mm. **B.** Fluorescent micrographs of 1:1 strain mixtures of indicated strains on TPM 1% agar at various times. Scale bar, 50 µm. Pearson’s correlation coefficients (right panels) for fluorescently labeled fruiting bodies formed from 1:1 strain mixtures at 24 h on TPM 1% agar. Value of 1 indicates perfect correlation between fluorescent channels, value of −1 indicates perfect segregation between fluorescent channels within a fruiting body. **C.** Stills from fluorescent timelapses of 1:1 strain mixtures in submerged culture at various times, scale bar, 50 μm **D.** Monocultures of WT or T6SS KO strains spotted next to one another on TPM agar after 1 week. Micrographs show colony interface. Arrows indicate chimeric fruiting bodies. Scale bars, 1000 µm and 100 µm.

Next, we used fluorescence microscopy to monitor cell dynamics during submerged culture development over 5 days. We found that despite vigorous initial mixing, in the mixture of the two wild type strains, the strains segregated within the first 24 h (Fig. 4C). As time progressed, the green A06 cells increased in dominance, while the red/magenta DK1622 population correspondingly decreased, indicating A06 antagonized DK1622 during development. The majority of fruiting bodies appeared around 72 h after induction of starvation and very few fruiting bodies formed from WT mixture compared to monoculture controls. Strikingly, fruiting bodies that formed fluoresced in only one channel, indicating they were composed of only one strain. To investigate the mechanism of developmental antagonism, we mixed the labeled T6SS mutants 1:1 and transferred them to starvation agar. At 24 h fruiting bodies were more numerous than WT mixtures. Some fruiting bodies were chimeric, composed of two strains that were generally stable. However, in other cases, strains segregated within their chimeric fruiting bodies over the course of the experiment (Fig. 4B).

To quantify the level of mixing within individual fruiting bodies, we measured fluorescent colocalization using Pearson’s correlation coefficient (Fig. 4B). For the fruiting bodies in the WT mixture, at 24 h most scored a Pearson’s correlation coefficient value of 0, resulting from only one fluorescence channel, i.e. clonal fruiting bodies. For the T6SS mutant mixtures, there was variability, but in general, these fruiting bodies exhibited a Pearson’s correlation coefficient value closer to +1, indicating some level of chimeric fruiting body formation. As time progressed, the Pearson’s correlation coefficient moved closer to −1, indicating more segregation as development progress (Fig. 4B). As a control, we mixed GFP and tdTomato labeled strains of DK1622 1:1. As expected, we observed Pearson’s correlation coefficient values close to +1 throughout the experiment, indicating sibling strains readily mixed and co-developed.

To understand strain interaction dynamics during development, we monitored 1:1 mixtures of the WT or T6SS mutants in submerged culture by time-lapse microscopy. For the WT mixtures, the strains segregated early, as they did on starvation agar (Fig. 4C). As time progressed, A06 gradually overtook DK1622, where nearly all fruiting bodies only consisted of A06, as determined by fluorescent microscopy. In the T6SS mutant mixtures, strains mixed and rippled for extended periods (days). At the end of the experiment, chimeric fruiting bodies formed, as they did on agar.

To imitate conditions more similar to wild conditions, we instead placed labeled cultures of WT or T6SS mutants next to one another on starvation agar and monitored interaction of the strains at the interface between colonies by fluorescence microscopy (Figure 4D). In the case of WT strains, the colony swarms did not merge. In contrast, with the T6SS mutants chimeric fruiting bodies at the colony interface were detected. We conclude that under starvation, A06 uses T6SS to antagonize and dominate DK1622. In the absence of T6SS, chimeric fruiting body formation occurred and strains within the chimera tend to segregate over time.

### Swarming behavior is altered in strain mixtures

To determine if the strains also segregate during vegetative growth, we spotted strain mixtures on rich media with either hard or soft agar, which promote either the A- or S-motility systems, respectively. On both hard and soft agar, monoculture controls of DK1622 or A06 with different fluorescent tags were well mixed (Fig. 5). Interesting, in mixtures of both WT and T6SS KO strains, swarm areas were reduced, indicating motility inhibition. In the WT mixtures, on hard agar little growth was apparent after one day, due to mutual antagonism. What patches of growth that were present, were segregated and strain specific. By three days, A06 had expanded and killed most of DK1622, except for some flares at the inoculum edge. On soft agar at one day, more cells were present than on hard agar, but growth was still reduced compared to monoculture controls. This implies that mutual killing occurred on soft agar, but was less efficient than on hard agar. Like on hard agar, patches of strains were segregated. Over the course of three days, the strains remained segregated and there were no obvious signs of antagonism. In the T6SS KO mixtures, on both hard and soft agar strains were mostly segregated at day one and this segregation persisted through the experiment.

**Fig. 5.**
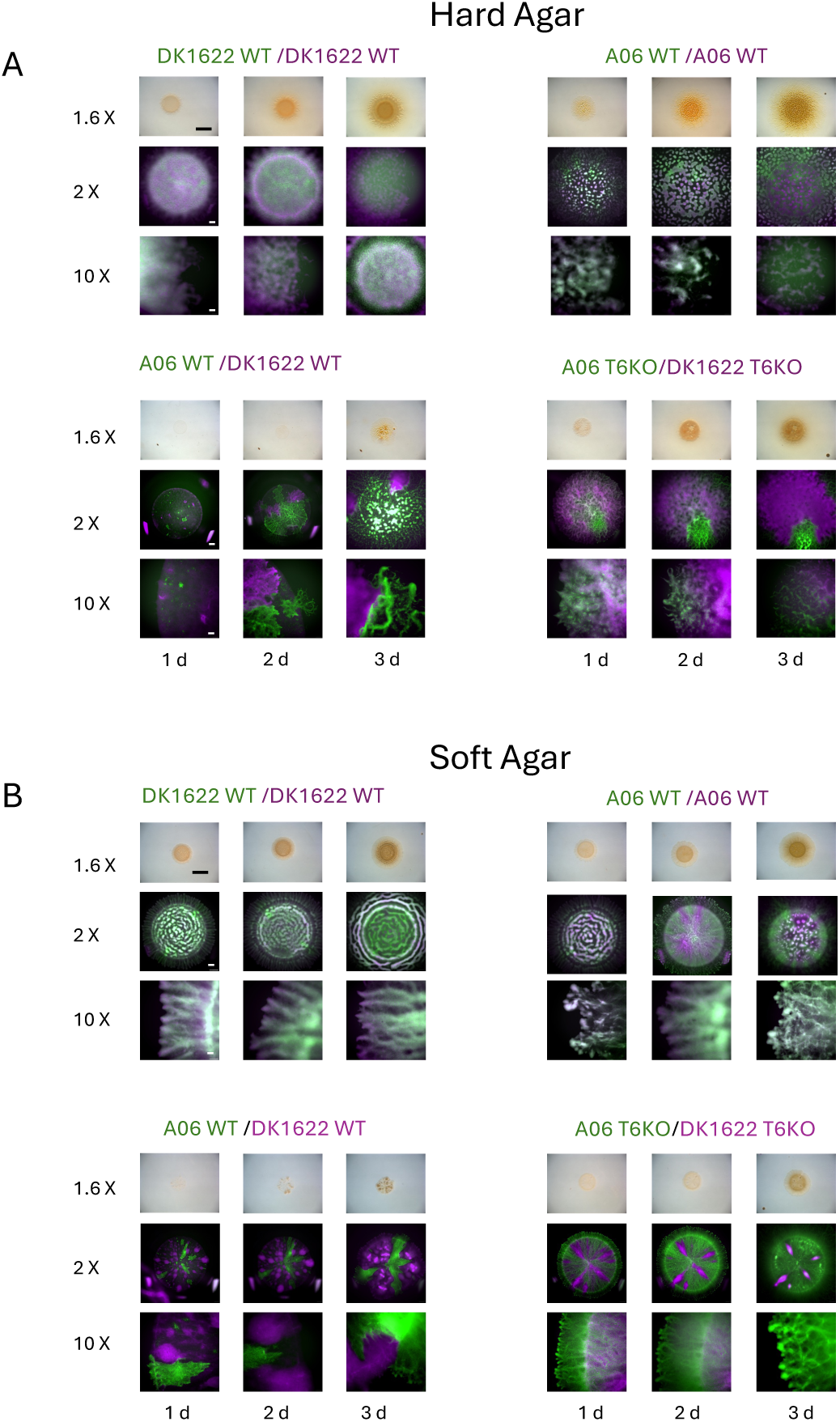
Swarming behavior of strain mixtures on hard and soft agar. **A**.1:1 strain mixtures spotted on CTT 1% agar. Images are of the same spot at different magnifications and at different time points. Scale bar for 1.6× = 5 mm, for 2× =500 μm, and for 10× =100 μm. **B**. Same as (A) but mixtures spotted on 0.5% agar.

### Sporulation efficiency reduced in WT mixtures but restored in T6SS mutant mixtures

Finally, we investigated sporulation efficiencies of WT and T6SS mutant mixtures spotted on starvation media, relative to their respective monocultures (Fig. 6). In WT mixtures, the sporulation efficiencies were lower than their respective monocultures, apparently caused by mutual antagonism. Importantly, in the T6SS mutant mixtures, sporulation levels returned to around monoculture levels, showing these divergent strains cooperated during development.

**Fig. 6.**
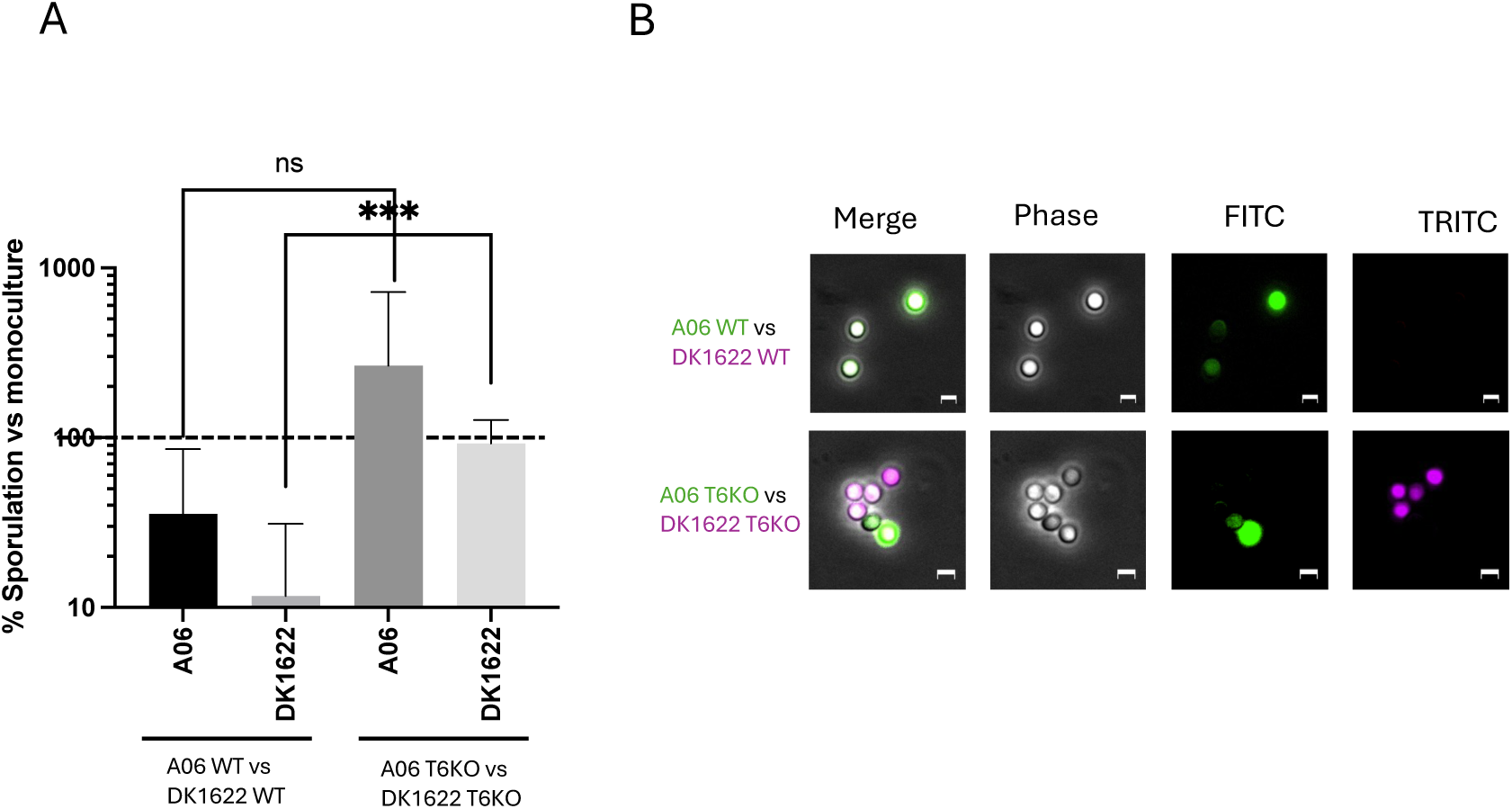
Sporulation compatibilities of strain mixtures. **A.** Sporulation efficiency of each strain in 1:1 mixtures relative to monoculture development after 5 days. ***P=0.0003. ns = not significant. **B.** Fluorescent micrographs of spores harvested from fruiting bodies after 1 week of development on TPM agar. Scale bars, 2 µm.

**Fig. 7.**
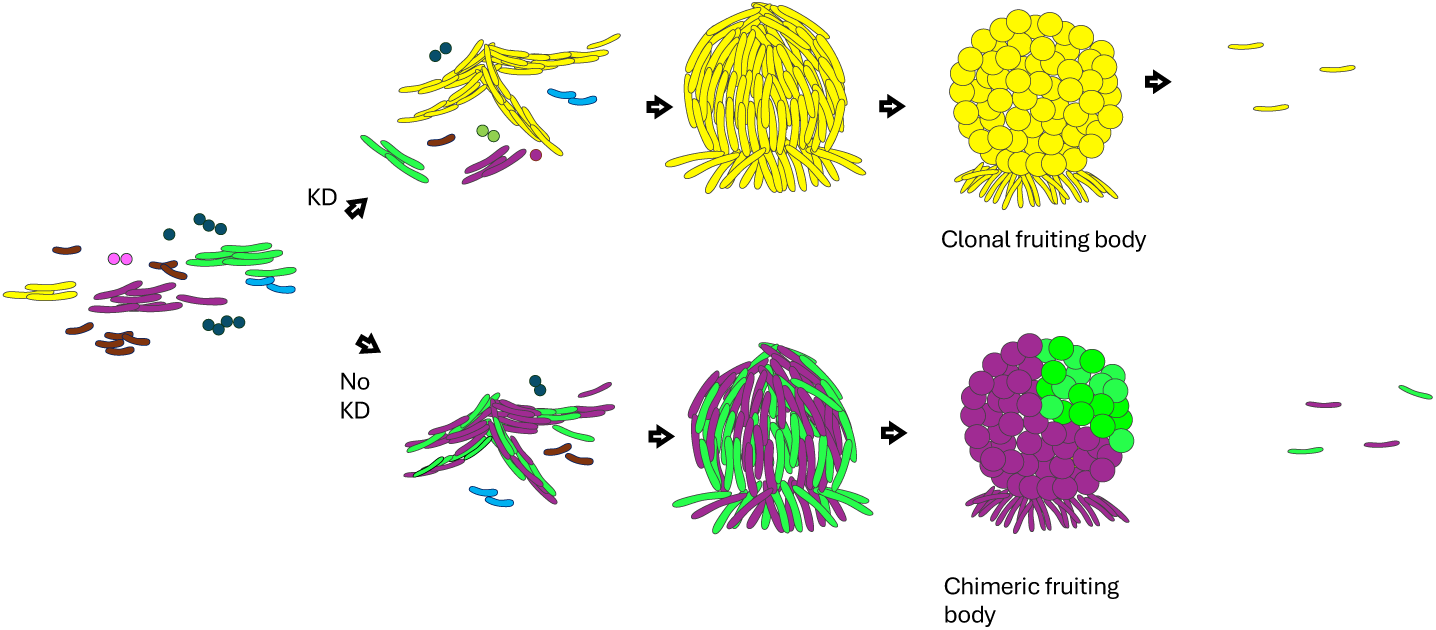
Model for the role of kin discrimination in fruiting body development. Rods represent vegetative myxobacteria cells; other shapes represent unrelated microbes from soil environments. Cells of the same color represent kin or clonal groups. When kin discrimination occurs (top), cells of one kin group aggregate to form a monoclonal fruiting body. Nonkin that attempt to join the aggregation are eliminated. In the absence of kin discrimination (bottom), distinct strains aggregate and form a chimeric fruiting body where subsequent segregation occurs.

## Discussion

### T6SS is a major determinant of kin discrimination between closely related strains

In a previous study, we found that OME, T6SS, and in some cases Rhs proteins, all function in kin discrimination between closely related strains [11]. Here, we report that T6SS is the dominant mechanism of kin recognition between distantly related *M. xanthus* strains. OME requires homotypic binding by two cells with compatible TraA receptors. TraA is highly polymorphic and although there is evidence of horizontal gene transfer of the *traAB* locus [11], the likelihood that distantly related strains contain compatible *traA* alleles is relatively low. The likely reason *traA* alleles are so divergent is because OME delivers polymorphic toxins, which creates selective pressure for OME to be highly specific and, hence, a high degree of diversity within *traA* alleles [8, 12]. Therefore, in many cases the KD function of OME is limited to closely related strains or strains that happen to have compatible receptors. Curiously, OME is not an ideal weapon to serve in KD, because it involves bidirectional toxin exchange so engaging in this behavior is often lethal between nonclonal cells. In addition, since *sitAI* gene cassettes are often associated with mobile genetic elements, it is likely that these mobile elements exploit OME to ensure their own retention and expansion in populations [33].

T6SS provides a broader mechanism for KD, since the delivery of toxins is unidirectional and lacks the specificity of OME. However, initiating contact still possesses the risk that the neighboring cell may return fire [34]. T6SS is knowns to function in intraspecific competition, targeting non-kin cells in other species, as found in *Vibrio* species [35–37] and *Serratia marcescens* [38]. In this study, we find that in *M. xanthus* strains with incompatible TraA receptors, T6SS is the major kin discriminating mechanism. However, the two strains have different genomic islands and genes, including *rhs* genes, so we cannot rule out that systems may play a small role, as well. Additionally, A06 contains 810 unique genes (with less than 50% identity) and DK1622 contains 690 unique genes, so some of these unique genes could contribute to antagonism.

We found that although there is mutual antagonism between strains, A06 uses its T6SS to dominate DK1622 under both vegetative and developmental conditions. There are several possible explanations for this result. The two strains may differ in their T6SS firing dynamics, with A06 deploying its T6SS faster or more frequently, or A06 may produce on average more T6SS complexes per cell. Other possibilities are A06 toxins are more potent or A06 has some cross-immunity toward DK1622 delivered toxins or A06 partially blocks toxin entry. Finally, since we observed that A06 swarms faster, it may be able to better evade incoming T6SS attacks and more quickly hunt down its targets.

### Chimeric development in T6SS KO mixtures

In mixtures of WT strains on starvation agar, results were similar to vegetative conditions − A06 dominated and nearly all fruiting bodies were composed entirely of A06. Strikingly, we found that in the absence of T6SS antagonism, chimeric fruiting bodies formed that produced viable spores from both strains. These chimeras were initially well-mixed, but over the course of several days, strains within fruiting bodies became segregated. The kin recognition mechanism causing this segregation is unknown but likely involves cell-surface associated molecule(s). The same mechanism that caused the strains to segregate during vegetative swarming could also promote segregation within fruiting bodies. Plausible candidate proteins involved in segregation are the divergent PilA and/or PilY1.1 proteins involved in type IV pili mediated (S-) motility.

Similar mixing experiments were done in the eukaryotic social amoeba *Dictyostelium discoideum*. Isolates of varying degrees of relatedness were mixed to determine their ability form chimeric fruiting bodies [39][40]. Here, adhesion between kin caused mixtures of unrelated strains to segregate within fruiting bodies, similar to our results in *M. xanthus*. Further, they found that the more distantly related two strains were, the greater the degree of segregation. Future studies in *M. xanthus* could similarly test whether strains with differing degrees of relatedness exhibit different degrees of segregation.

In WT mixtures, the sporulation of both strains was reduced, due to antagonism. In T6SS KO mixtures, A06 sporulated more efficiently in a mixture than in a monoculture. These findings are consistent with other studies that found some strains sporulate more efficiently in mixtures than in monocultures due to unknown synergistic or exploitation effects [41–43]. However, the sporulation of DK1622 was nearly the same in the T6SS KO mixture as in monoculture, indicating that any synergistic effects benefited only A06. One candidate to explain this synergy could be C-signal. The C-signal is cell-surface localized, and its transmission is contact dependent [44, 45]. During development, critical concentrations of the C-signal must be reached for aggregation and then sporulation to occur. The receptor for the C-signal is unknown, but once sufficient C-signal is present, gene expression is altered to initiate sporulation [46]. The C-signal is a product of the *csgA* gene, which is 98.8% identical between the two strains, so presumably the same C-signal is functional on both strains. A06 may be more sensitive to the C-signal, either by expressing more C-signal receptors or by requiring a lower concentration of C-signal to alter gene expression.

### Strain segregation during swarming

Both WT strain mixtures and T6SS KO mixtures segregated on hard and soft agar within the first 24 hours. The mechanism of segregation is unknown. It is possible that the strains produce a surface molecule that preferentially recognizes and binds to itself. Such candidates include PilA or PilY1.1, since again these strains contain very different alleles for these genes. Furthermore, studies on *Vibrio* found that cells expressing the same allele of pilin aggregate together, causing them to segregate from strains expressing a different allele [34, 47]. This has been proposed as a mechanism that allows cells to aggregate and defend themselves from rival T6SS attacks [34]. Strain segregation may be maintained by the ‘corpse barrier effect,’ where dead cells in the boundary between the two strains prevent further strain mixing [34, 48].

We found mixtures of both WT or T6SS mutants had reduced swarming on both hard and soft agar compared to monocultures. For the WT mixture, this can be explained, at least in part, by the drastic reduction in cell number due to mutual antagonisms. Notably, T6SS KO mixtures had increased swarming compared to WT mixtures, but still less than monocultures. This might be caused by reduction in cell number due to antagonism by Rhs proteins or other mechanisms. The polymorphic nature of PilA/PilY1.1 between these strains could also contribute to reduced swarm expansion. In other words, social motility may be more efficient when swarms have the same PilA pilin and PilY1.1 tip adhesion types. This is plausible given that *Vibrio* strains expressing the same *pilA* gene auto-aggregate [49]. To address these possibilities, future studies could employ single cell tracking and *pilA*/*pilY1.1* allele swaps between strains.

TraA is a cell surface receptor that is normally present at low levels, but when *traAB* is overexpressed, TraA functions as an adhesin and causes cells expressing the same *traA* allele to adhere [31, 50]. Future work could investigate the effect of overexpression of *traAB* in these strains. If the native *traAB* in each strain was overexpressed, this would likely make the segregation more dramatic. However, if two strains were engineered to overexpress the same *traA* allele, it may cause the two strains to adhere, reducing the amount of segregation between T6SS KO strains.

## Acknowledgements

This work was supported by the National Institutes of Health grants R35GM140886 to D.W.

